# Osteocyte Perilacunar/canalicular Remodeling (PLR) Drives Spatially Heterogeneous Lacunar Remodeling During and After Lactation

**DOI:** 10.64898/2026.06.01.729140

**Authors:** Xiaoyu Xu, Yilu Zhou, Wonsae Lee, Srijani Datta, Joel Boerckel, Liyun Wang, X. Sherry Liu

## Abstract

**Background:** Pregnancy and lactation impose substantial demands on maternal calcium homeostasis, leading to pronounced skeletal remodeling during lactation followed by recovery after weaning. Although bone mass is largely returned at the tissue level after weaning, it remains unclear whether osteocyte-level remodeling exhibits a similarly reversible pattern. Osteocytes regulate mineral mobilization through perilacunar/canalicular remodeling (PLR), which is elevated during lactation. However, its spatial and temporal regulation in response to reproduction remains poorly defined.

**Objective/Hypothesis:** This study aims to determine whether PLR-regulated lacunar remodeling during reproduction varies with osteocyte location and relative age. We hypothesized that osteocyte PLR-mediated lacunar change is spatially heterogeneous during lactation, varies with osteocyte location and relative age, and may persist after weaning.

**Methods:** Female wild-type (WT) and osteocyte-specific PTH/PTHrP receptor knockout (cKO) mice were studied across reproductive stages. Longitudinal *in vivo* DXA and μCT were used to assess skeletal changes. Sequential fluorochrome labeling tracked mineral dynamics and defined osteocyte relative age. Osteocyte PLR activity was evaluated by MMP13 immunohistochemistry. Lacunar-canalicular structure (LCS) was assessed using Ploton silver nitrate staining, and spatially resolved lacunar morphology was quantified using high-resolution backscattered scanning electron microscopy (bSEM).

**Results:** At the tissue level, reproduction induced distinct skeletal responses, characterized by reversible cortical bone loss and persistent trabecular deterioration. Cortical bone loss during lactation was spatially asymmetric and confined to the posterior cortex. Fluorochrome labeling further resolved surface-specific remodeling patterns during reproduction, including endocortical resorption at the posterior cortex and sustained deposition at anterior endocortical and posterior periosteal surfaces.

At the cellular level, osteocyte PLR activity increased during lactation in WT mice and returned after weaning, whereas no changes were observed in cKO mice. Consistently, lacunar size increased during lactation and returned toward baseline after weaning in WT mice but remained unchanged in cKO mice.

Spatially resolved analysis demonstrated that lacunar remodeling was heterogeneous across cortical thickness. At the anterior cortex, lacunar enlargement occurred near the endocortical surface during lactation and was reversible after weaning. In contrast, at the posterior cortex, lacunar enlargement occurred near the periosteal surface and persisted after weaning. These spatial patterns corresponded to cortical regions enriched with newly formed osteocytes, whereas pre-existing osteocytes exhibited minimal changes. This spatial heterogeneity was absent in cKO mice.

**Conclusion:** Osteocyte PLR-mediated lacunar remodeling during reproduction is spatially heterogeneous and varies with osteocyte location and relative age. These findings demonstrate that recovery at the tissue level does not necessarily extend to the osteocyte microenvironment and identify osteocyte PLR-mediated lacunar remodeling as a spatially structured and potentially persistent component of reproductive skeletal adaptation. Together, these results highlight a previously unrecognized layer of maternal skeletal health.

## 1. Introduction

Pregnancy and lactation impose substantial calcium demands on the maternal skeleton to support fetal development and milk production [1,2]. During lactation, bone undergoes marked mineral loss and structural remodeling, followed by substantial recovery after weaning [1,3,4]. Although this process has traditionally been considered reversible at the whole-bone level, growing evidence suggests that reproduction may leave lasting alterations in skeletal microstructure [4–8]. These findings raise the possibility that recovery of bone mass after weaning may not fully reflect restoration of bone quality at the microstructural or tissue levels.

Osteocytes are central regulators of bone mass, mineral homeostasis, and skeletal adaptation in response to hormonal and mechanical cues [9–11]. Embedded within the mineralized matrix, osteocytes form an extensive lacunar-canalicular system (LCS) that supports mechanosensing, intercellular communication, and regulation of bone remodeling [12–16].

Through perilacunar/canalicular remodeling (PLR), osteocytes modify their surrounding matrix by resorbing and replacing perilacunar mineral [12,17,18]. This process alters lacunar morphology and local matrix properties, enabling mineral mobilization during periods of increased physiological demand [2,12,18] and thereby influencing bone tissue quality.

During lactation, osteocyte PLR is activated in response to elevated calcium demand, and enlargement of osteocyte lacunae has been reported during this period [4,17,19,20]. These changes are mediated, at least in part, through PTH/PTHrP receptor (PPR) signaling in osteocytes, as osteocyte-specific deletion of PPR suppressed PLR activity during lactation [19]. However, whether lactation-induced lacunar enlargement fully resolves following weaning remains incompletely explained. In particular, although bone mass is largely returned after weaning, it is not known whether the osteocyte-level mineral remodeling follows a similarly reversible trajectory.

In addition to temporal regulation, osteocyte behavior is shaped by spatial and cellular context within cortical bone [14,20,21]. The organization of osteocyte lacunae contributes to local tissue material properties, and changes in lacunar size or LCS connectivity may influence osteocyte mechanotransduction and intercellular signaling [12,13,16,20,22]. Thus, recovery of bone mass and cortical geometry after weaning does not necessarily indicate restoration of the osteocyte-level mechanical or structural environment. Moreover, osteocyte morphology and PLR activity vary across cortical regions [14,20,21], reflecting differences in local remodeling dynamics and mineralization history.

Beyond spatial variation, osteocytes also differ in relative age according to the timing of their incorporation into the matrix [10,23–27]. Newly formed osteocytes, embedded during periods of active bone formation, may experience distinct microenvironments and behave differently compared to pre-existing osteocytes formed before reproduction. These differences raise the possibility that osteocytes of different ages may respond differently to the metabolic demands of lactation.

Taken together, we hypothesize that osteocyte PLR-mediated lacunar alterations during reproduction is spatially heterogeneous and depends on both osteocyte location and relative age. Using longitudinal *in vivo* imaging, sequential fluorochrome labeling, and high-resolution backscattered scanning electron microscopy, we will define the temporal-spatial dynamics of lacunar alterations throughout reproduction. To further establish the role of osteocyte PLR in mediating these changes, we will compare osteocyte-specific PPR knockout mice, in which lactation-induced PLR is abolished [19], with wild-type littermate controls.

## 2. Method

### 2.1 Animals and Experimental Design

All animal procedures were approved by the Institutional Animal Care and Use Committee (IACUC #804378) at the University of Pennsylvania. Eight-week-old female osteocyte-specific PTH/PTHrP Receptor (PPR) conditional knockout mice (cKO; 14kb-Dmp1-Cre^+/-^; PPR^fl/fl^) and their littermate wild-type controls (WT; PPR^fl/fl^) were used in this study [19,28]. Although dentin matrix protein (DMP1) is also expressed in late-stage osteoblasts, these cells are considered to be destined to differentiate into osteocytes [26]. Genotyping was performed using standard polymerase chain reaction (PCR) protocols as detailed in Supplementary Materials (Table S1).

Two complementary experimental designs were employed: 1) Longitudinal Study (Fig. 1A): WT and cKO mice were randomly assigned to Virgin and Reproduction (Repro) groups (n = 3-4 per group). Mice in the Repro group were paired at 8 weeks old and followed through the reproductive cycle, including pregnancy (21 days), lactation (12 days), and post-weaning recovery (14 days). Animals were sacrificed at 15 weeks of age following the recovery period. 2) Cross-sectional Study: WT and cKO mice were randomly assigned to three groups: Virgin, Lactating (Lac), and Post-weaning (Wean, n = 4-6 per group). Mice in the Lac and Wean groups were mated at 10 and 8 weeks old, respectively, and underwent pregnancy (21 days) followed by 12 days of lactation. Mice in the Wean group were allowed to recover for 14 days after weaning. All animals were euthanized at 15 weeks of age at different reproductive stages, along with age-matched Virgin controls. For both longitudinal and cross-sectional studies, litter sizes were normalized to 6 pups per dam within 48 hours after birth to ensure consistent lactational demand.

**Fig. 1.**
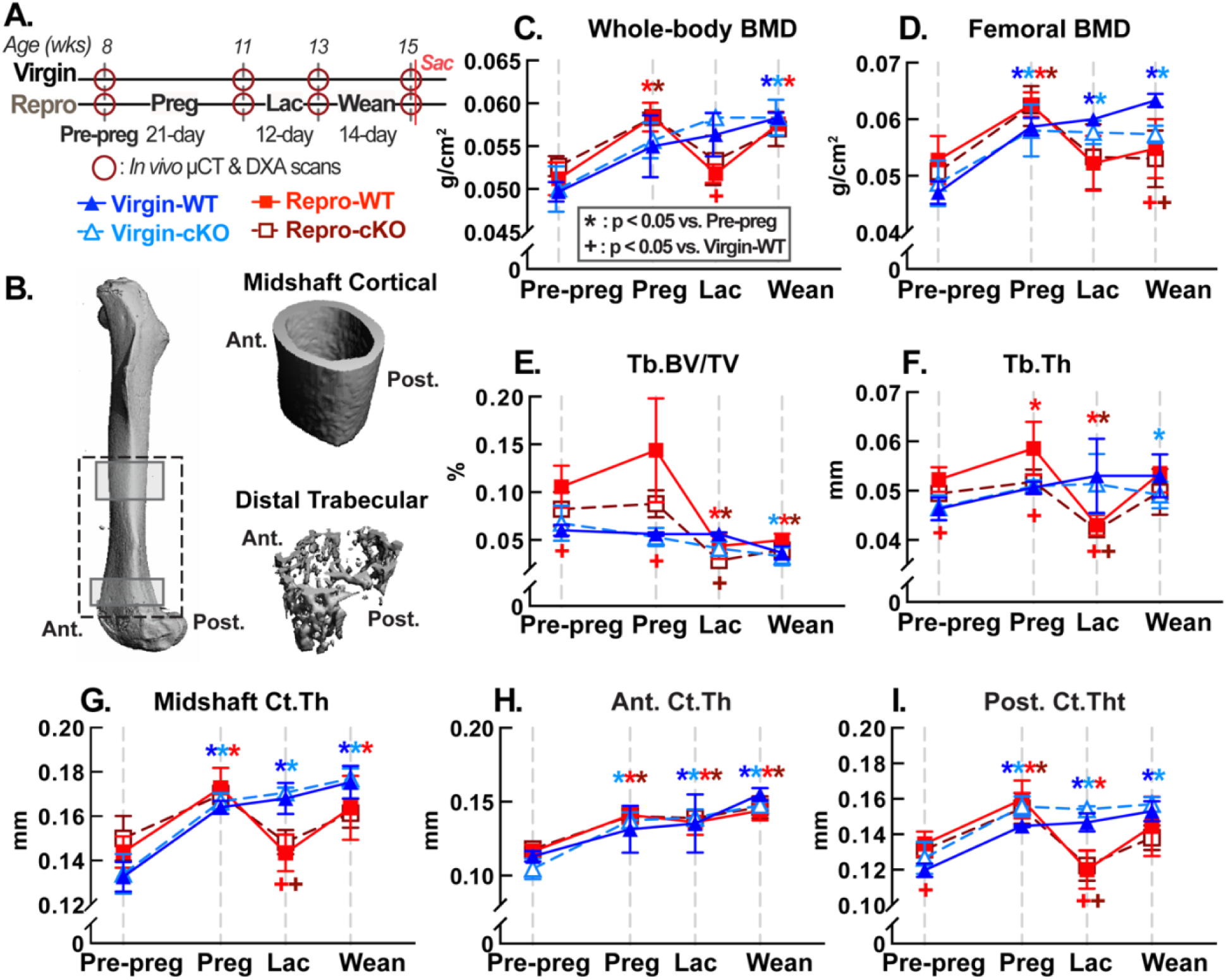
Longitudinal analysis of bone mass and skeletal morphology in reproductive (Repro) and age-matched virgin (Virgin) female mice with osteocyte-specific PPR deletion (cKO) and littermate controls (WT) (A) Experimental timeline showing reproduction strategy and *in vivo* imaging schedule. (B) μCT regions of interest at the distal trabecular and midshaft cortical femur; anterior (Ant.) and posterior (Post.) cortices were analyzed separately. Whole-body and femoral BMD were measured by DXA (C, D). Trabecular BV/TV (E) and Tb.Th (F), and cortical thickness at the midshaft (G), anterior (H), and posterior cortices (I), were quantified by μCT. Differences relative to the pre-pregnancy baseline were assessed within each group using paired Student’s t-tests (*p < 0.05). Differences relative to Virgin-WT at each time point were assessed by one-way ANOVA (**+**p < 0.05).

All animals were maintained under a 12-hour light/dark cycle with ad libitum access to food (Calcium: 0.81%, Phosphorus: 0.61%, and magnesium: 0.21%; 5053 PicoLab Rodent Diet 20, LabDiet, Richmond, IN, USA) and water. Virgin mice were group-housed (three animals per cage) under standard conditions. Reproductive mice were individually housed during the final week of pregnancy and throughout lactation to avoid stress-induced infanticide but group-housed during post-weaning recovery.

### 2.2 *In vivo* imaging of the dual-energy X-ray absorption (DXA) and micro-computed tomography (μCT) scans

To longitudinally assess skeletal and mineral changes during reproduction, *in vivo* μCT and DXA scans were performed on animals in the longitudinal cohort at four time points (Fig. 1A): pre-pregnancy (pre-preg; 8 weeks old), post-labor (Preg; 11 weeks old), end of the lactation (Lac; 13 weeks old), and the end of post-weaning (Wean; 15 weeks old). Virgin mice were scanned at corresponding time points.

*In vivo* μCT scans (Scanco vivact80, Scanco Medical AG, Brüttisellen, Switzerland) were performed on the left femur, which was fixed in a customized holder to minimize motion during the scan [29]. Scans were acquired over an 8.5 mm-length region of interest (ROI) spanning the midshaft-to-distal femur, positioned proximal to the distal growth plate (Fig. 1B). Scanning was conducted at a voxel size of 14 μm with 55 kVp, 145 μA, and 200 ms integration time. The total scan time was approximately 40 min per animal. DXA scans were performed using an iNSiGHT VET DXA (Scintica, Webster, TX, USA) following the same schedule as μCT imaging. Each animal underwent three consecutive whole-body scans (excluding skull), each lasting approximately 30 secs, using settings of 60 kVp (low energy), 80 kVp (high energy), 0.8 mA, and 5-second integration time. Final measurements were averaged across the three scans at each time point. All animals were anesthetized with isoflurane (2.5% induction/maintenance) during *in vivo* imaging and allowed to recover under a heating lamp after scanning.

DXA-derived parameters included the areal bone mineral density (aBMD, g/cm^2^) for whole-body (excluding skull; Whole-body BMD) and the right femora (Femoral BMD). For trabecular bone analysis by μCT, a 60-slice (0.84 mm) volume of interest (VOI) was identified in the distal femoral metaphysis (Fig. 1B), beginning proximal to the growth plate and excluding the primary spongiosa and cortical shell [11,30]. VOIs were identified by visual inspection to ensure consistency across animals [31]. Trabecular bone was manually contoured and segmented using a global threshold of 417 mg HA/cm^3^ after Gaussian filtering (sigma = 0.8, support = 1) [32]. Standard trabecular parameters included bone volume fraction (BV/TV), trabecular thickness (Tb.Th), trabecular number (Tb.N), trabecular spacing (Tb.Sp), connectivity density (Conn.D), and structure model index (SMI). For cortical bone analysis, an 85-slice (approximately 1.2 mm) VOI was identified at the femoral midshaft (Fig. 1B) and segmented semi-automatically using manufacturer-provided software (Scanco Medical AG). Images were filtered (Gaussian filter, sigma = 0.8, support = 1) and thresholded at 584 mg HA/cm^3^. Cortical parameters included cortical area (Ct. Area), cortical thickness (Ct.Th), and polar moment of inertia (pMOI). Cortical thickness at the anterior (Ant. Ct.Th) and posterior (Post. Ct.Th) cortices was also quantified to assess the site-specific changes.

### 2.3 Multi-fluorochrome Labeling and Multiplexed cryohistology

Fluorochrome labeling was performed on WT and cKO mice in the Reproduction (Repro) group of the longitudinal cohort to track mineralization dynamics across reproductive stages. A sequence of multicolor fluorochrome labels was administered to mark bone formation at distinct time points (Fig. 2A), including a subcutaneous injection of calcein green (G, 15 mg/kg) at the onset of pregnancy (8 weeks of age), an intraperitoneal injection of alizarin red complexone (R, 30 mg/kg) at the onset of lactation (11 weeks of age), and an intraperitoneal injection of tetracycline yellow (Y, 30 mg/kg) at the onset of post-weaning (13 weeks of age). Virgin mice were subjected to the same labeling schedule as the Repro group. All animals were euthanized at 15 weeks of age.

**Fig. 2.**
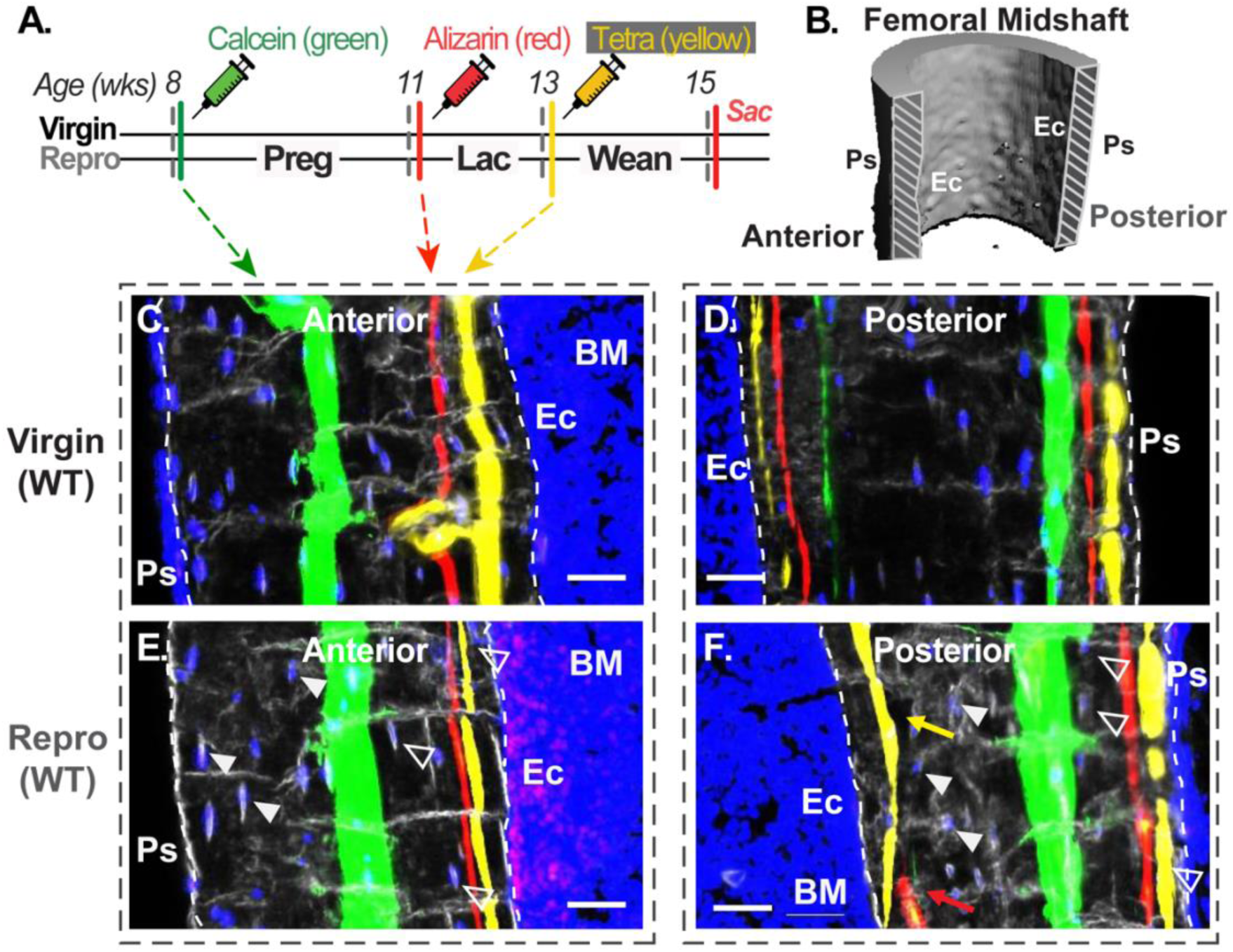
Fluorochrome labeling of dynamic mineral remodeling at the anterior and posterior cortices of the femoral midshaft in reproductive (Repro) and age-matched virgin (Virgin) mice. (A) Experimental timeline and fluorochrome labeling strategy. (B) Representative 3D reconstruction showing full cortical thickness from periosteal (Ps) to endosteal (Ec) surfaces at the femoral midshaft. Grey-shaded regions indicate the anterior and posterior cortical areas analyzed. (C-F) Fluorochrome labeling at anterior (C, E) and posterior (D, F) cortices in Virgin-WT and Repro-WT mice. White dashed lines indicate cortical boundaries. Solid arrows (E, F) indicate pre-existing osteocytes formed prior to reproduction, whereas open arrows indicate osteocytes formed during reproduction. In (F), the red arrow indicates loss of pre-lactation mineral labeling, and the yellow arrow indicates mineral deposition during weaning. Similar spatial mineral remodeling patterns were observed between WT and cKO mice at both anterior and posterior cortices in Virgin and Repro groups (Fig. S2). Fluorescence labels: calcein (green), alizarin (red), tetracycline (Tetra; yellow), and To-PRO-3 (blue; nuclei). BM: bone marrow cavity. Scale bar: 30 μm.

Following euthanasia, the right femur was harvested and fixed in 4% paraformaldehyde (PFA) at 4 °C for 48 hours, subsequently rinsed in 1x phosphate-buffered saline (PBS), and cryoprotected in a graded sucrose solution (10% followed by 20%) for a total of 48 hours.

Samples were then embedded in Tissue-Tek O.C.T. Compound (Sakura Finetek USA Inc., Torrance, CA, USA) using liquid nitrogen. Undecalcified longitudinal cryo-sections (8 μm thickness), spanning the midshaft to distal region with both anterior and posterior cortices exposed, were prepared using Cryofilm IIC tape (Section-Lab Co., Hiroshima, Japan) as previously described [33,34]. Cryo-sections were mounted onto glass slides using 1% chitosan adhesive, dried at 4 °C for 48 hours, rehydrated in 1x PBS for 15 minutes, and cover-slipped with a nuclear counterstaining solution (1:1000 dilution of TO-PRO-3 iodide in 50% glycerol in 1x PBS; Invitrogen, ThermoFisher Scientific, USA).

Mounted cryo-sections were scanned at 10x magnification using an Axio Scan Z1 (Zeiss, Germany). Both dark-field and fluorescent images were acquired as described previously [9], using appropriate filters to detect the calcein (FITC), alizarin (RFP), tetracycline (customized filter [34]), and nuclear (Cy5) signals.

Longitudinal cryo-sections enabled visualization of the full cortical thickness, spanning from the endocortical surface (Es; adjacent to the bone marrow cavity) to the periosteal surface (Ps; outer cortical boundary, Fig. S2). This approach allows spatial assessment of mineral deposition and remodeling patterns across the entire anterior and posterior cortices at the femoral midshaft.

### 2.4 Immunohistochemistry (IHC) and Assessment of Osteocyte PLR Activity

To evaluate osteocyte-mediated perilacunar/canalicular remodeling (PLR) activity and validate the efficiency of the PPR deletion in osteocytes, the expression of matrix metalloproteinase-13 (MMP13), a well-established marker of PLR activity [19,33,35], was evaluated by immunohistochemistry (IHC) in cortical bone of the tibial midshaft in Virgin, Lactating (Lac), and Post-weaning (Wean) groups of both WT and cKO mice from the cross-sectional study cohort (Fig. 3A).

**Fig. 3.**
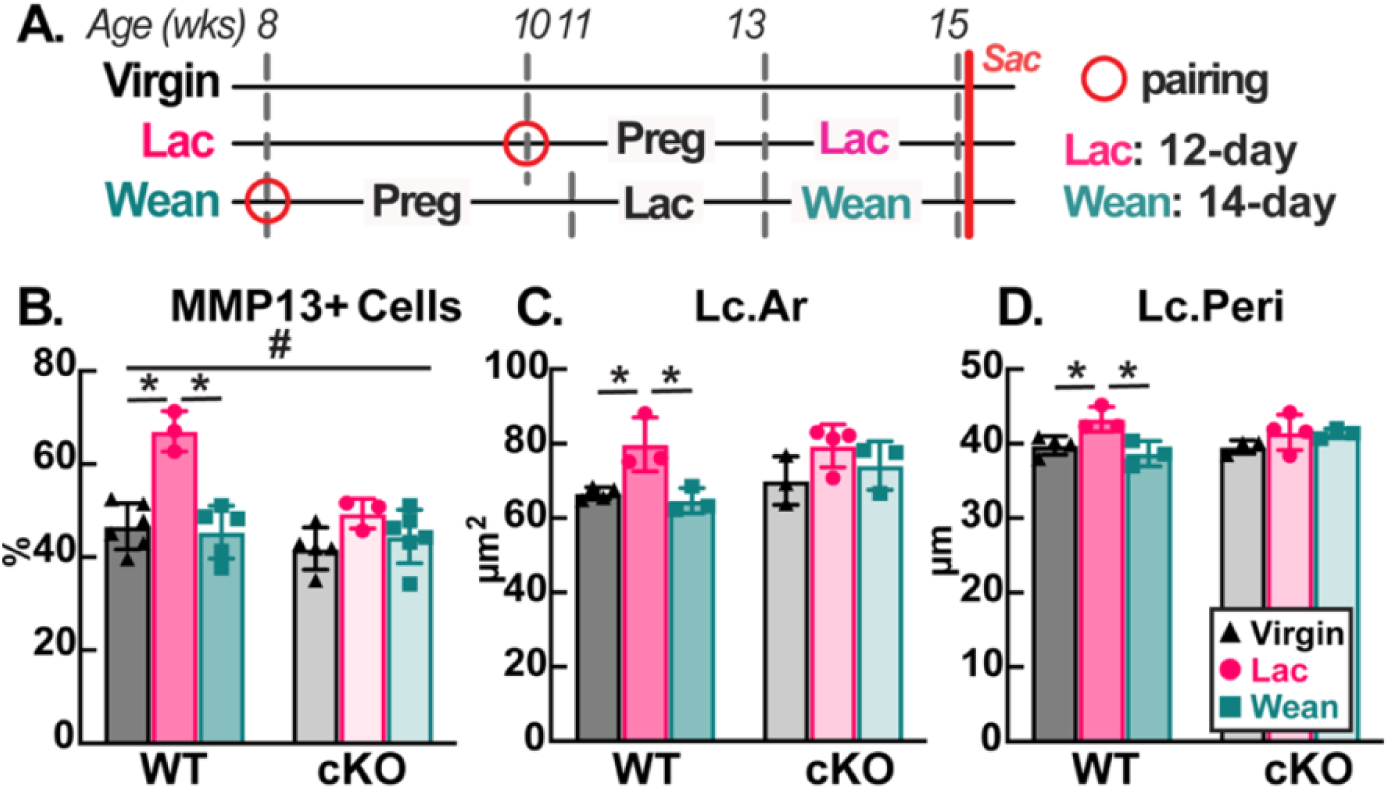
Osteocyte PLR activity and lacunar morphology in WT and cKO mice across reproductive stages. Reproductive groups included lactation (Lac), post-weaning (Wean), and age-matched virgin controls (Virgin). (A) Experimental timeline. (B) Percentage of MMP13-positive osteocytes (MMP13+) assessed by immunohistochemistry (IHC). Lacunar area (Lc.Ar, C) and perimeter (Lc.Peri, D) were measured by Ploton’s silver nitrate staining. Differences among Virgin, Lac, and Wean groups were analyzed separately in WT and cKO mice using one-way ANOVA (*p < 0.05). The effects of reproductive status and genotype, and their interaction, were assessed using two-way ANOVA, with significant interactions indicated by # (p < 0.05).

Following euthanasia, left tibiae were harvested and fixed in 4% PFA at 4 °C for 48 hours, followed by decalcification in 10% ethylenediaminetetraacetic acid (10% EDTA; pH 7.4) at 4 °C for 3 weeks. Decalcified samples were dehydrated through a graded ethanol series, embedded in paraffin, and sectioned at 7 μm thickness at the midshaft region with lateral and medial cortices exposed.

Paraffin sections were deparaffinized, subjected to antigen retrieval, and incubated with a primary antibody against MMP13 (1:150; Abcam, ab39012). MMP13 immunoreactivity was detected using a biotinylated secondary antibody, followed by avidin-conjugated peroxidase and chromogenic substrate (Innovex Universal Animal IHC kit), as previously described [33,35]. Sections were counterstained with methyl green (1%; R&D Systems, Minneapolis, NM, USA) for nuclear visualization and mounted for imaging. Negative controls were prepared using the rabbit serum IgG.

Bright-field images were acquired at 20x magnification using the Axio Scan Z1 (Zeiss, Germany) and analyzed by ImageJ/Fiji (NIH) [36]. For each section, a 0.5 mm-length region of interest (ROI) was defined at the tibial midshaft for both medial and lateral diaphyses. Within each ROI, osteocytes were identified based on lacunar morphology and the presence of nuclear staining (methyl green-positive). Lacunae containing a stained nucleus were counted and defined as the total osteocyte population. Lacunae exhibiting both nuclear staining and detectable cytoplasmic MMP13 signal were classified as MMP13-positive osteocytes (Fig. S1). The percentage of MMP13-positive osteocytes (MMP13+ Ocys; %) was calculated as the ratio of MMP13-positive osteocytes to the total number of osteocytes. For each animal, values were averaged across medial and lateral ROIs for each section (n=3 sections per animal).

**2.5 Ploton Silver Nitrate Staining for Lacunar-Canalicular Structure (LCS) Quantification**

Ploton silver nitrate staining was performed on a separate set of paraffin sections (7 μm thickness) obtained from the tibial midshaft of mice in the cross-sectional cohort, as described in Section 2.4. Following previous established protocols [4,33,37], sections were deparaffinized and rehydrated, then incubated in a freshly prepared silver nitrate solution for 60 minutes in the dark. Sections were subsequently rinsed in 5% sodium thiosulfate for 10 minutes, dehydrated, and mounted. Stained sections were imaged using bright-field microscopy under the same imaging conditions described in Section 2.4. Lacunae were identified based on their characteristic morphology and enhanced contrast provided by silver deposition [33,37]. For each section, lacunae were randomly selected from both medial and lateral cortices (n=15 per cortex side; 30 lacunae total per section), within the same ROIs defined in Section 2.4. Quantitative measurements, including lacunar area (Lc.Ar), perimeter (Lc.Peri), and the number of canaliculi per lacuna (canaliculi # per lacuna), were manually obtained using OsteoMeasure software (OsteoMetrics, Inc., Decatur, GA, USA). For each animal, measurements were first averaged across three sections. Group-level values were then calculated by averaging across animals within each reproductive group for both WT and cKO cohorts.

### 2.6 Backscatter Scanning Electronic Microscopy (bSEM) and Osteocyte Lacunar Quantification

To evaluate changes in osteocyte lacunar morphology during and after lactation, lacunae within the femoral diaphyseal cortex were analyzed through bSEM for the cross-sectional cohort (Fig. 3A).

Following euthanasia, left femora were harvested, cleaned of surrounding soft tissue, and processed for polymethyl methacrylate (PMMA) embedding, as previously described [19,20].

Embedded specimens were longitudinally sectioned through the midshaft-to-distal region along the medial-lateral axis using an IsoMet low-speed saw equipped with a diamond wafering blade (102 x 0.3 mm; Buehler, IL, USA). Specimens were divided into equal halves, exposing both anterior and posterior cortical surfaces. Exposed surfaces were sequentially polished using carborundum papers of decreasing grit size (1 μm and 0.3 μm; 3M Lapping Discs, Ted Pella Inc., CA, USA), followed by fine polishing at 0.06 μm colloidial silica suspension (Ted Pella, Inc.), as described previously [19,20]. Samples were then carbon-coated to enable high-resolution bSEM imaging (Cell and Developmental Biology Microscopy Core, University of Pennsylvania, PA, USA). bSEM images were acquired from both anterior and posterior cortices at the femoral midshaft using an FEI Quanta 250 FEG (ThermoFisher; 15 kV accelerating voltage, 9 mm working distance) at a resolution of 1209 pixels/mm.

Images were thresholded, and osteocyte lacunae were identified based on their characteristic morphology. Lacunar area (Lc.Ar) was quantified using a customized MATLAB (MathWorks, Natick, MA, USA) analysis program. Lacunae intersecting image borders were excluded from analysis. To assess site-specific differences in lacunar dimension, lacunar measurements were first grouped by anatomical location (anterior vs. posterior) and analyzed across the full cortical thickness, spanning from the endocortical to the periosteal surfaces. For each location, approximately 140-160 lacunae were quantified per sample. To further evaluate spatial heterogeneity, cortex at each location was subdivided into five equal sub-regions along the periosteal-to-endocortical axis (0-100%; Fig. S3). With each sub-region, 20-30 lacunae were analyzed, enabling region-specific quantification of lacunar morphology. This spatially resolved approach allowed quantitative assessment and direct comparison of regional changes in lacunar dimension between reproductive groups and genotypes.

### 2.7 Statistical Analysis

All data are presented as mean ± standard deviation (SD) unless otherwise specified. Assumptions of normality and homogeneity of variance were tested using the Shapiro-Wilk test and Levene’s test, respectively, prior to statistical analyses. Statistical analyses were performed using GraphPad Prism (GraphPad Software, San Diego, CA, USA).

For the longitudinal study, differences in skeletal morphology across time points within the same group (Virgin or Repro) were assessed relative to the pre-pregnancy baseline (Pre-preg) using paired Student’s *t*-tests. At each corresponding time point, comparisons to the Virgin-WT group were conducted using one-way analysis of variance (ANOVA) with Bonferroni post hoc correction. For the cross-sectional study, differences in osteocyte-related measurements, including MMP13 expression by IHC and LCS parameters by the silver nitrate staining, among Virgin, Lactation, and Post-weaning groups were analyzed separately within WT and cKO cohorts using one-way ANOVA, followed by Bonferroni post hoc correction for multiple comparisons. The interactive effect of genotype and reproductive group on the MMP13 expression and LCS parameters was assessed using two-way ANOVA. For spatially resolved analyses by bSEM, differences in lacunar morphology among Virgin, Lactation, and Post-weaning groups were performed within corresponding sub-region using one-way ANOVA with Bonferroni post hoc correction, separately for anterior and posterior cortices in WT and cKO cohorts.

A value of p < 0.05 was considered statistically significant. Statistical significance is indicated in figures as follows: p < 0.05 (*), p < 0.01 (**), and p < 0.001 (***). Values of p < 0.1 are reported as trends, with the exact *p*-value provided.

## 3. Results

### 3.1 Osteocyte-specific PPR deletion suppresses osteocyte PLR without altering baseline skeletal and reproductive phenotypes

Deletion of PPR in osteocytes was first confirmed by PCR detection of transgenic DNA segments for PPR and Cre recombinase, followed by gel electrophoresis. To functionally validate osteocyte-specific PPR deletion in osteocytes, MMP13 expression was assessed as a functional readout of osteocyte-mediated perilacunar/canalicular remodeling (PLR) activity (Fig. S1). In WT mice, the percentage of MMP13-positive osteocytes increased during lactation by approximately 44% compared to the age-matched Virgin controls (Fig. S1([A-Z]), B). In contrast, cKO mice showed no detectable change in MMP13 expression during lactation relative to age-matched Virgin mice (Fig. S1([A-Z]), D).

Baseline physiological characteristics were comparable between genotypes. Virgin mice exhibited continuous weight gain during normal growth, with no differences observed between WT and cKO groups (Table S2([A-Z])). Across reproduction, body weight followed the expected physiological pattern. In WT mice, body weight increased markedly during pregnancy (+30%) and lactation (+44%), both significantly higher than age-matched Virgin mice, and decreased after weaning (−16%) (Table S2([A-Z])). No genotype-dependent differences were observed at any time point. Reproductive performance was also similar between genotypes, with litter size of 6-7 pups per dam at parturition, normalized to 6 pups per dam within 48 hours after birth (Table S2([A-Z])). All litters were maintained successfully during the lactation period, with no evidence of filial cannibalism or offspring abandonment. Average pup body weight at weaning was 5.6 ± 0.5 g for the WT mice and 4.9 ± 0.3 g for the cKO mice (Table S2([A-Z])).

Together, these results confirm that the conditional knockout model used in this study achieves effective functional suppression of PLR activity, without affecting baseline growth or reproductive performance. This validates the osteocyte-PPR deletion model for investigating the role of osteocyte PLR during reproduction.

### 3.2 Reproduction induces dynamic skeletal changes with reversible cortical but persistent trabecular bone loss

In Virgin mice, bone mass increased progressively with age, with continuous increases in whole-body and femoral aBMD and no differences between genotypes (Fig. 1C, D). In contrast, mice in the Repro group exhibited dynamic changes across reproductive stages (Fig. 1C, D). Whole-body and femoral aBMD increased during pregnancy (+14% and +19%, respectively), decreased during lactation (−11% and −17% vs. Preg), and recovered after waning (+11% and +5% vs. Lac), reaching levels comparable to age-matched Virgin controls at 15 weeks of age (Fig. 1C, D).

Bone microstructure was further evaluated using the *in vivo* μCT. In Virgin mice, trabecular bone at the proximal femur remained stable with age in WT but showed age-related decline in cKO mice, as demonstrated by a reduction in BV/TV (−51% at Wean vs. Pre-preg; Fig. 1E), accompanied by decreased Tb.N and increased Tb.Sp (Table S3([A-Z]),B). In contrast, Repro mice exhibited pronounced trabecular alterations across reproductive stages. In WT mice, trabecular bone mass increased during pregnancy, with BV/TV and Tb.Th elevated by 36% and 12%, respectively (Fig. 1E, F). During lactation, trabecular deterioration was observed, with marked reductions in BV/TV (−59%), Tb.Th (−17%), Tb.N (−24%), and Conn. D (−73%), accompanied by increases in Tb.Sp (+33%) and SMI (1.4-fold) (Fig. 1E, F; Table S3). These changes persisted after weaning, with only Tb.Th returning to pre-pregnancy levels comparable to the age-matched Virgin controls (Fig. 1F). Similar changes were also observed in cKO mice across reproductive stages (Fig. 1E-G; Table S4). These changes indicate persistent trabecular bone loss and long-lasting structural deterioration following reproduction regardless of genotype.

In contrast, cortical bone at the femoral midshaft exhibited a distinct remodeling pattern. Virgin animals showed continuous cortical expansion during growth in both WT and cKO mice, whereas Repro mice displayed dynamic but reversible changes (Fig. 1G). In WT mice, Ct.Th, Ct.Area, and pMOI increased during pregnancy (+20%, +20%, and +21%, respectively, vs. Pre-preg), decreased during lactation (−15%, −17%, and −10%, respectively, vs. Preg), and fully recovered after weaning (Fig. 1G; Table S4). By the end of the post-weaning period, Ct.Th, Ct.Area, and pMOI were returned to levels comparable to age-matched Virgin controls and remained 14%, 15%, and 19%, respectively, above the pre-pregnancy baseline (Fig. 1G; Table S4). In cKO mice, changes in Ct.Th across reproductive stages followed a temporal trend similar to that in WT mice (Fig. 1G), whereas Ct.Area and pMOI remained unchanged throughout reproduction (Table S4).

Together, the DXA and μCT analyses demonstrated that the magnitude and temporal pattern of changes in bone mineral density, trabecular microarchitecture, and cortical geometry were generally comparable between genotypes. Cortical and trabecular compartments exhibited distinct structural responses to reproduction regardless of genotype: cortical bone fully recovered after weaning, whereas lactation-induced microstructural deterioration in trabecular bone persisted throughout the post-weaning period.

### 3.3 Cortical remodeling during lactation is spatially asymmetric and localized to the posterior cortex

The cortical bone was further evaluated separately at the anterior and posterior sites. In WT mice, Virgin animals exhibited continuous cortical expansion at both anterior and posterior sites, with greater Ct.Th at anterior (+37%) compared to the posterior site (+28%) by 15 weeks of age (Fig. 1H, I). In contrast, Repro mice displayed site-dependent cortical remodeling across reproductive stages. Posterior cortical thickness increased during pregnancy (+19% vs. Pre-preg), decreased during lactation (−25% vs. Preg), and fully recovered after weaning (Fig. 1I). By the end of the post-weaning period, posterior Ct.Th was 9% higher than the pre-pregnancy baseline and comparable to age-matched Virgin controls (Fig. 1I). At the anterior site, Ct.Th increased during pregnancy (+21% vs. Pre-preg) and remained stable during lactation (+17% vs. Pre-preg) (Fig. 1H). After weaning, anterior Ct.Th continued to increase (+24% vs. Pre-preg) but remained 7% lower than that of age-matched Virgin controls (Fig. 1H, I).

No genotype-dependent differences were observed in the magnitude or temporal pattern of cortical thickness changes at either the anterior or posterior cortex across reproductive stages. These results showed that bone loss at the femoral midshaft during lactation was confined to the posterior cortex, whereas the anterior cortex maintained continuous growth across reproduction.

### 3.4 Fluorochrome labeling shows spatially distinct mineral remodeling between cortical endocortical and periosteal surfaces during reproduction

By employing sequential fluorochrome labeling at pre-pregnancy, end of pregnancy, and end of lactation, we further visualized dynamic mineralization patterns across the cortical thickness, extending from the endocortical surface to the periosteal surface at the anterior and posterior cortices of the femoral midshaft using longitudinal cryo-sections (Fig. 2A, B; Fig. S3).

Consistent with the μCT results, Virgin WT mice exhibited continuous mineralization during growth. At the posterior cortex, symmetric G-R-Y labeling was present at both endocortical and periosteal surfaces (Fig. 2D), whereas at the anterior cortex, this sequential labeling was primarily localized to the endocortical surface, with minimal activity at the periosteal surface (Fig. 2C).

In Repro WT mice, the anterior cortex showed a mineralization pattern similar to that in Virgin mice (Fig. 2E). Although no mineral resorption was detected during lactation, fluorochrome label spacing during pregnancy and lactation (G-R labels) was reduced relative to age-matched Virgin mice (Fig. 2C, E). Together with μCT results, mineral remodeling at the anterior cortex during reproduction was spatially dependent, with deposition confined primarily to the endocortical surface and minimal activity at the periosteal surface (Fig. 2E).

In contrast, the posterior cortex exhibited a distinct spatial-dependent pattern. At the endocortical surface, mineral resorption occurred during lactation, evidenced by the loss of G-R fluorochrome labels compared to Virgin mice (Fig. 2D, F). This pattern was consistent with the reduced Ct.Th measured at the posterior cortex by μCT (Fig. 1I). Following weaning, mineralization resumed at the endocortical surface, as shown by the appearance of the yellow fluorochrome label (Fig. 2F). In contrast, mineral deposition persisted at the periosteal surface (Fig. 2F), with sequential G-R-Y labels incorporation from pregnancy through post-weaning, following a pattern similar to that in Virgin mice (Fig. 2D, F). Together with the μCT data, bone loss at the posterior cortex during lactation was localized to mineral resorption at the endocortical surface.

These spatial patterns were observed in both WT and cKO mice, with no detectable differences between genotypes (Fig. S2). The fluorochrome labeling patterns were consistent with μCT findings of site-specific bone loss at the posterior cortex during lactation and showed spatially dependent remodeling between endocortical and periosteal surfaces at the femoral midshaft.

### 3.5 Sequential Fluorochrome labeling delineates osteocyte relative age across cortical regions

Based on the timing of fluorochrome administration, labeled regions corresponded to bone formed during defined reproductive stages, including pregnancy (G-R), lactation (R-Y), and post-weaning (Y- surface) (Fig. 2E, F; Fig. S2([A-Z])).

Using this temporal framework, osteocytes were categorized according to their relative formation time. Osteocytes embedded in unlabeled bone regions corresponded to bone formed prior to pregnancy and were classified as ‘pre-existing osteocytes’. In contrast, osteocytes located within labeled regions were associated with bone formed during or after pregnancy and were classified as ‘newly formed osteocytes’.

Across the cortical regions, these osteocyte populations exhibited distinct spatial distributions. Pre-existing osteocytes were primarily located in cortical regions formed prior to reproduction, including the central cortex at the posterior site and the periosteal-to-central region at the anterior site, where mineral remodeling was relatively restricted (Fig. 2E, F). In contrast, newly formed osteocytes were enriched in regions undergoing active bone remodeling during reproduction, particularly near the anterior endocortical surface and both endocortical and periosteal surfaces at the posterior cortex (Fig. 2E, F).

This spatial distribution provided a framework to distinguish osteocyte populations based on their relative formation time and exposure to reproductive stages, enabling subsequent evaluation of lacunar remodeling in relation to osteocyte location and relative age.

### 3.6 Lactation induces osteocyte PLR activity and lacunar structural remodeling that are abolished in osteocyte-specific PPR-deficient mice

In WT mice, the percentage of MMP13-positive osteocytes increased to 67% in lactating mice compared to 47% in age-matched Virgin controls and decreased to 45% after weaning, returning toward the baseline level of Virgin mice (Fig. 3A). In contrast, cKO mice exhibited no detectable change in MMP13 expression, with values remaining approximately at 47%, across reproductive stages (Fig. 3A).

The LCS morphology was further assessed using Ploton silver nitrate staining. In WT mice, lacunar size increased during lactation, with lacunar area (Lc.Ar) and perimeter (Lc.Peri) elevated by +20% and +9%, respectively, compared to the age-matched Virgin controls (Fig. 3B, C). Following weaning, lacunar dimensions decreased (Lc.Ar: −18% and Lc.Peri: −11% vs. Lac), returning to levels comparable to age-matched Virgin mice (Fig. 3B, C). In cKO mice, no changes in lacunar dimensions were observed across reproductive stages, with Lc.Ar and Lc.Peri remaining at approximately 75 μm^2^ and 41 μm, respectively (Fig. 3B, C). Across all measurements, LCS parameters in cKO mice were comparable among Virgin, Lactation, and Post-weaning groups (Fig. 3B, C).

Together, changes in MMP13 expression and lacunar dimensions were observed in WT mice during lactation and returned after weaning, whereas these changes were not observed in cKO mice.

### 3.7 Spatially heterogeneous lacunar remodeling varies with osteocyte location and relative age and is absent in osteocyte-specific PPR-deficient mice

Lacunar morphology across the cortical thickness was evaluated at the anterior and posterior sites of the femoral midshaft using high-resolution bSEM. For each site, osteocyte lacunae were analyzed within five sub-regions defined along the periosteal-to-endocortical axis to characterize spatial variation of lacunar remodeling during reproduction (Fig. S3).

In WT mice, lacunar remodeling exhibited pronounced spatial heterogeneity across the cortical thickness during reproduction. At the posterior cortex, lacunar size remained stable in regions spanning from the endocortical surface to the mid-cortex during lactation (Fig. 5F). In contrast, lacunae located closer to the periosteal surface (0-40%) enlarged during lactation (Lc.Ar: +28% and +34% vs. Virgin for 0-20% and 20-40% regions, respectively) and remained elevated after weaning (+23% and +31% vs. Virgin), showing persistent enlargement across the post-weaning period (Fig. 5G). At the anterior cortex, a different spatial pattern was observed during reproduction. Lacunar size remained largely unchanged in regions spanning from the periosteal surface to the mid-cortex (Fig. 4F). In contrast, lacunae located near the endocortical surface (80-100%) enlarged during lactation (+34% vs. Virgin) and returned toward baseline levels after weaning (−13% vs. Lac), reaching values comparable to those of age-matched Virgin controls (Fig. 4G).

**Fig. 4.**
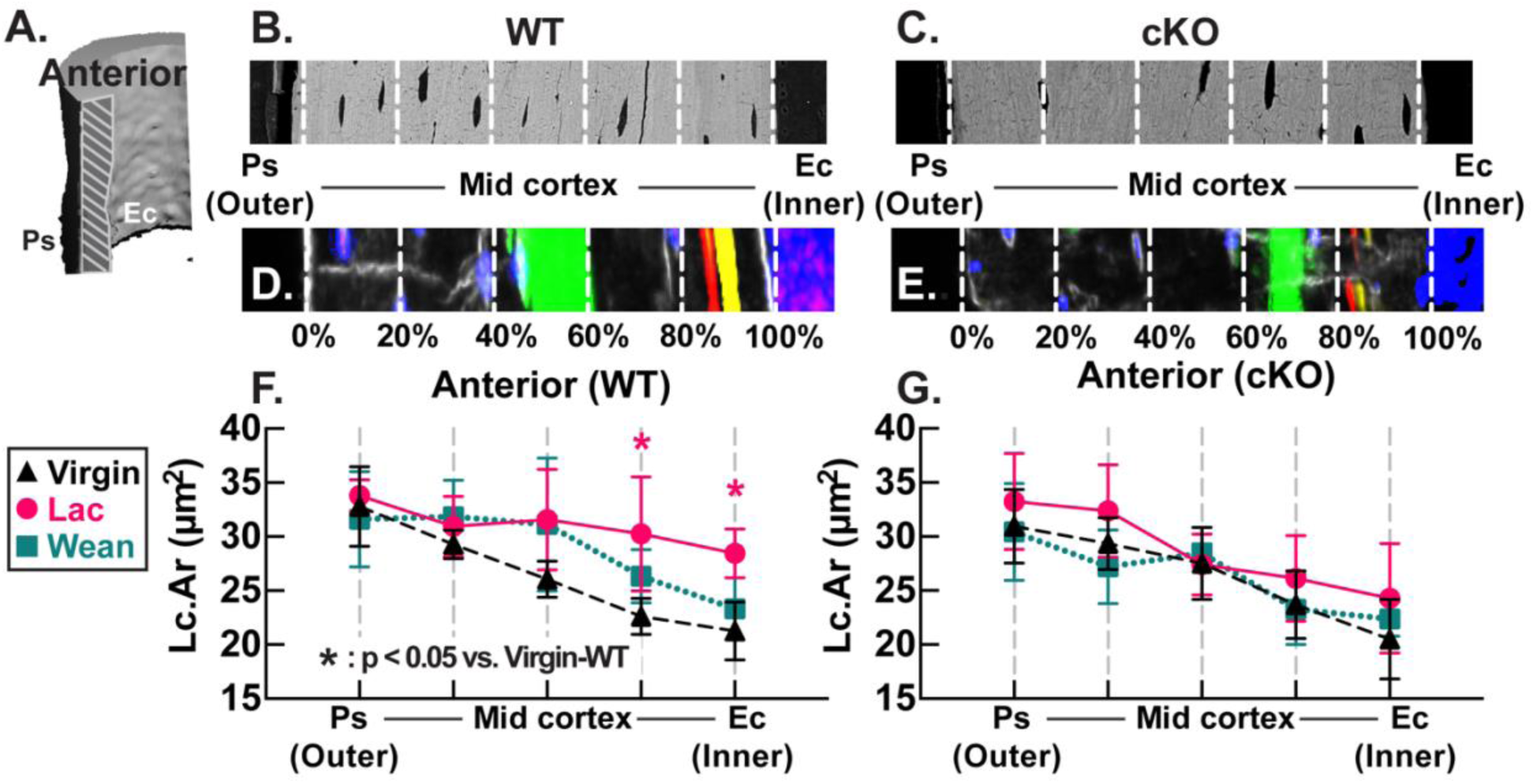
Spatially resolved analysis of lacunar morphology at the anterior cortex of the femoral midshaft. (A) Representative 3D reconstruction of the longitudinal region of interest (ROI) spanning the full cortical thickness from periosteal (Ps) to endosteal (Ec) surfaces for bSEM imaging. (B, C) Representative bSEM images of the anterior cortex in post-weaning WT (B) and cKO (C) mice, showing five equal sub-regions across the cortical thickness (0-100% from Ps to Ec; white dashed lines). (D, E) Representative fluorochrome labeling images during reproduction in WT (D) and cKO (E) mice (adapted from Fig. 3 and Fig. S2). (F, G) Spatially resolved quantification of lacunar area (Lc.Ar) across cortical sub-regions in Virgin, Lactation (Lac), and Post-weaning (Wean) groups of WT (F) and cKO (G) mice. Differences among groups were assessed within each sub-region using one-way ANOVA (*p < 0.05).

**Fig. 5.**
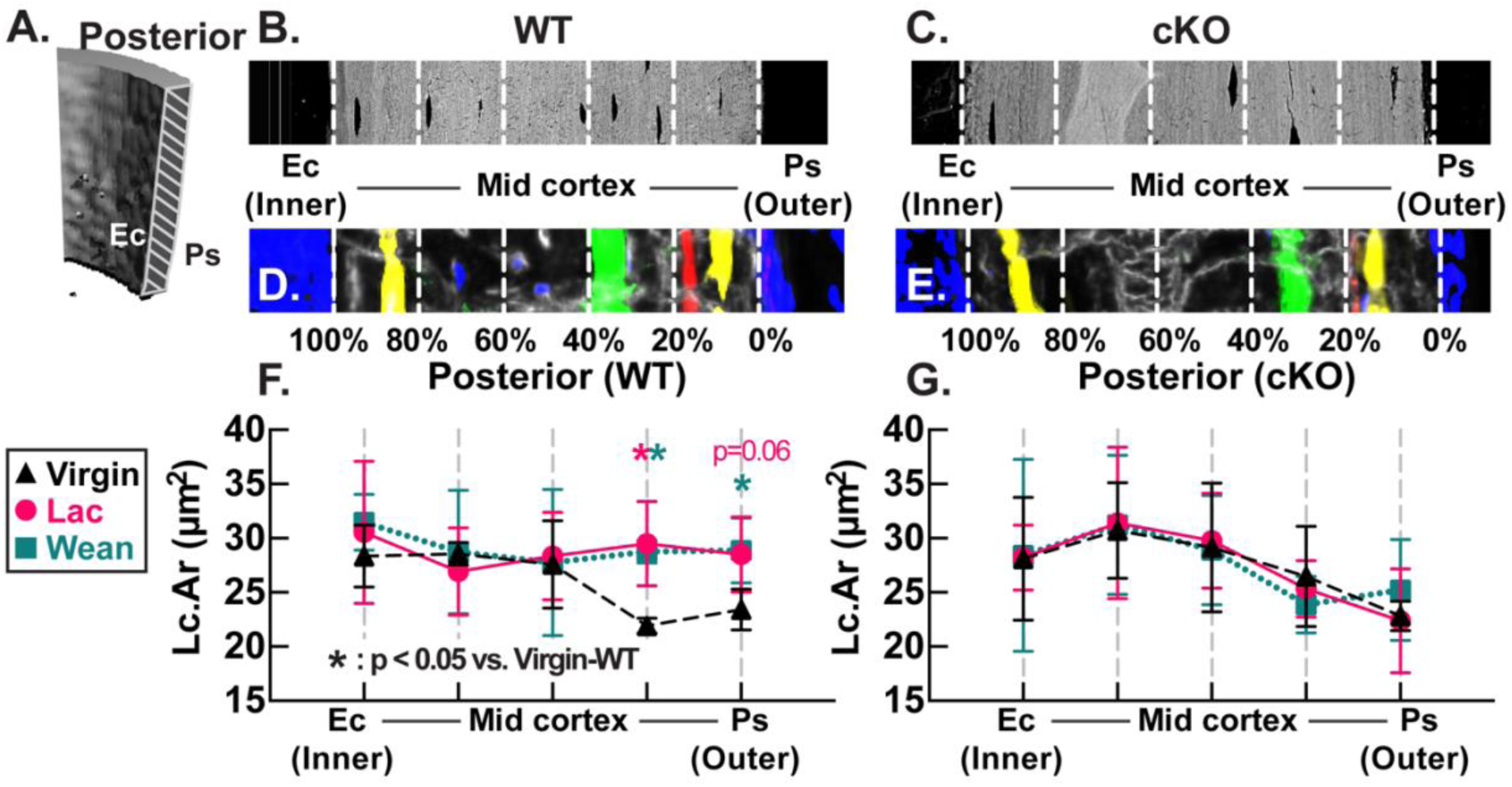
Spatially resolved analysis of lacunar morphology at the posterior cortex of the femoral midshaft. (A) Representative 3D reconstruction of the longitudinal region of interest (ROI) spanning the full cortical thickness from periosteal (Ps) to endosteal (Ec) surfaces for bSEM imaging. (B, C) Representative bSEM images of the posterior cortex in post-weaning WT (B) and cKO (C) mice, showing five equal sub-regions across the cortical thickness (0-100% from Ps to Ec; white dashed lines). (D, E) Representative fluorochrome labeling images during reproduction in WT (D) and cKO (E) mice (adapted from Fig. 3 and Fig. S2). (F, G) Spatially resolved quantification of lacunar area (Lc.Ar) across cortical sub-regions in Virgin, Lactation (Lac), and Post-weaning (Wean) groups of WT (F) and cKO (G) mice. Differences among groups were assessed within each sub-region using one-way ANOVA (*p < 0.05). Trends are indicated for 0.05 ≤ p < 0.1.

Across both anterior and posterior sites, lacunar size varied systematically along the cortical thickness. Stable lacunar size was observed in regions with minimal cortical mineral deposition, extending from mid-cortex to periosteal (anterior) or endocortical (posterior) surface (Fig. 4F; Fig. 5F). In contrast, dynamic lacunar remodeling was localized to cortical surfaces undergoing active mineral deposition, with reversible enlargement at the endocortical surface (anterior) and persistent alteration at the periosteal surface (posterior) (Fig. 4G; Fig. 5G). Furthermore, when mapped to osteocyte populations defined by mineral labeling, regions exhibiting dynamic lacunar remodeling corresponded to locations enriched with ‘newly formed osteocytes’ (Fig. 4D, F; Fig. 5D, F), whereas regions with stable lacunar size corresponded to locations enriched with ‘pre-existing osteocytes’ (Fig. 4D, F; Fig. 5D, F).

In cKO mice, this spatial variation in lacunar remodeling was not observed at either anterior or posterior sites (Fig. 4G; Fig. 5G). With the suppression of osteocyte PLR activity during lactation, lacunar size remained unchanged throughout reproduction across all sub-regions corresponding to pre-existing or newly formed osteocytes (Fig. 4E, G; Fig. 5E, G).

## 4. Discussion

This study investigated the spatial heterogeneity of osteocyte PLR-mediated lacunar remodeling in the maternal skeleton during reproduction. A *Dmp1*-Cre-induced osteocyte-specific PPR conditional knockout mouse model (cKO), together with littermate controls (WT), were used, combining with longitudinal *in vivo* imaging, sequential fluorochrome labeling, immunohistochemistry, and high-resolution backscattered SEM (bSEM), to resolve lacunar remodeling patterns across maternal cortical bone.

To define tissue-level skeletal changes across reproduction, bone mass and structural parameters were assessed longitudinally by *in vivo* DXA and μCT. Virgin WT and cKO mice showed similar age-related growth, indicating that osteocyte-specific PPR deletion did not affect baseline skeletal development. During reproduction, lactation induces substantial bone loss in WT mice, followed by recovery after weaning. However, trabecular and cortical compartments exhibited distinct responses. Trabecular bone at the distal femur showed marked deterioration during lactation with incomplete post-weaning recovery, indices of bone volume fraction and connectivity remained impaired. In contrast, cortical bone at the femoral midshaft exhibited a largely reversible response. These findings are consistent with prior rodent studies demonstrating reversible cortical bone loss but persistent trabecular bone structural deterioration following reproduction [7,17,19–21]. Notably, pregnancy induced minimal trabecular changes in both WT and cKO mice, differing from prior rat studies reporting pregnancy-associated trabecular bone loss [1,4,7,8,38], potentially due to species-specific differences [2,39,40]. Overall, these bone mass and structural changes during reproduction were comparable between WT and cKO mice, suggesting that osteocyte-specific PPR deletion does not substantially alter global bone mass or skeletal structure during reproduction.

Although μCT analysis indicates overall recovery of cortical bone after weaning, site-specific analyses of the anterior and posterior cortices revealed a spatially asymmetric pattern across the femoral cortex. Lactation-induced bone loss was localized primarily to the posterior cortex, whereas the anterior cortex continued to expand throughout reproduction. Sequential fluorochrome labeling further localized these changes to specific bone surfaces. In Virgin mice, mineral deposition followed distinct regional patterns, with labeling present at both endocortical and periosteal surfaces in the posterior cortex but primarily restricted to the endocortical surface in the anterior cortex. This spatial asymmetry may partly reflect cortical drift during radial growth together with regional differences in local mechanical strain distribution [14,41–43].

Mechanical loading in mouse lone bones generates spatially heterogeneous cortical strain, with compressive and tensile forces distributed across opposite cortices [30,44–46]. At the femoral diaphysis, the anterior cortex is predominately subjected to compressive strain, whereas the posterior cortex experiences tensile strain [14]. Accordingly, local mechanical stimulation may be greater at anterior endocortical and posterior periosteal surfaces, potentially promoting site-specific mineral deposition during cortical growth [46,47].

During reproduction, the posterior cortex exhibited a distinct remodeling response. Loss of pre-lactation fluorochrome labels at the posterior endocortical surface indicated active mineral resorption during lactation, followed by renewed mineral deposition after weaning. In contrast, mineral deposition at the posterior periosteal cortex persisted throughout reproduction, maintaining a pattern similar to the anterior endocortical surface. Together, these results demonstrate that cortical remodeling during reproduction is highly spatially organized by both anatomical location and bone surface, with lactation-induced mineral resorption confined to the posterior endocortical surface. Notably, this spatial remodeling pattern was preserved in both WT and cKO mice.

Previous studies in rat and mouse models have reported increased LCS remodeling and lacunar enlargement in maternal cortical bone during lactation [4,17,20,21]. However, many of these analyses treated osteocyte lacunae across the cortex as a single population, averaging lacunar changes over entire cortical regions, which may obscure local differences in osteocyte behavior [17,20,21]. Osteocytes are derived from late-stage osteoblasts and interact dynamically with the surrounding mineral matrix during embedding [9,10], and their activity is strongly influenced by the local microenvironment [11,23,48,49]. During bone remodeling, osteocytes located near active resorption/formation surfaces may be exposed to different biochemical and mechanical conditions compared to those embedded in regions with minimal turnover. In addition to spatial location, osteocyte activity also varies with the relative age [9,48,50,51], which is associated with their differentiation stage (early stage vs. mature) and defined by the timing of incorporation into the mineral matrix. Osteocytes generated during pregnancy, lactation, or post-weaning are typically located near active mineralizing surfaces, whereas osteocytes formed before reproduction are predominantly found in the central cortex. These spatial and temporal differences suggest that osteocyte responses to reproductive demands may not be uniform across the cortex. In our study, the anterior and posterior cortices were therefore subdivided into five regions along the endocortical-to-periosteal axis to better evaluate lacunar remodeling with respect to osteocyte location and relative age. In WT mice, lacunar remodeling during reproduction followed a region-specific pattern that paralleled mineral remodeling but did not fully reflect tissue-level recovery. Lacunar enlargement during lactation occurred preferentially in regions undergoing active mineralization, particularly near the anterior endocortical and posterior periosteal surfaces. The extent of recovery after weaning further varied between cortical regions. At the anterior cortex, lacunar enlargement near the endocortical surface was transient during lactation and returned toward baseline after weaning. In contrast, at the posterior cortex, lacunar enlargement near the periosteal surface persisted beyond weaning.

By comparison, lacunae located in regions with minimal mineralization, specifically the anterior mid-to-periosteal and posterior mid-to-endocortical regions, remained relatively stable throughout reproduction. Therefore, despite full recovery of cortical bone mass and geometry after weaning at the whole bone level, osteocyte-level lacunar remodeling remains spatially heterogeneous. Cortical regions exhibiting dynamic changes in lacunar size were enriched with osteocytes formed during reproduction, whereas regions containing pre-existing osteocytes formed prior to reproduction showed minimal lacunar changes. This age-dependent osteocyte response provides a mechanistic basis for the spatial heterogeneity of lacunar remodeling.

To delineate the role of osteocyte PLR in mediating these spatially dependent lacunar changes, we further examined osteocyte-specific PPR knockout mice. In cKO mice, the lactation-induced increase in MMP13 expression and accompanying lacunar enlargement observed in WT mice were absent. Moreover, the spatial heterogeneity of lacunar remodeling was no longer evident, as lacunar size remained relatively stable across cortical regions and reproductive stages. These findings indicate that osteocyte PLR is required not only for the lactation-induced lacunar remodeling but also for establishing the spatially organized and age-dependent osteocyte responses within cortical bone. This observation extends previous studies linking PPR signaling to lactation-induced PLR and further suggests that osteocyte PTH/PTHrP signaling regulates the age-dependent osteocyte responses that underlie the heterogeneous spatial distribution of lacunar remodeling across the cortex [1,19,52].

This spatially heterogeneous osteocyte remodeling may have important implications for bone quality and mechanical behavior of maternal skeletons. Lacunar porosity and LCS connectivity contribute to cortical porosity and tissue-level material properties [12,20,53,54], and region-specific changes in lacunar size may lead to localized differences in stiffness, strain distribution, and interstitial fluid flow [13,55–57]. Thus, even when cortical bone mass recovers after weaning, persistent alterations in lacunar morphology may influence osteocyte mechanosensitivity and long-term mechanical adaptation of the maternal skeleton. Therefore, osteocyte-level remodeling represents an additional level of skeletal adaptation that is not reflected in conventional measurements of bone mass or structure.

While our results strongly suggest that osteocyte-level remodeling during reproduction may influence bone quality beyond what is captured by tissue-level measurements, several limitations should be considered when interpreting these findings. First, although spatial and temporal patterns of lacunar remodeling were clearly defined, the direct mechanical consequences of these lacunar changes were not evaluated. As a result, the effects of region-specific lacunar alterations on local material properties and tissue-level mechanical behavior remain unclear. Future studies integrating nanoindentation, Raman spectroscopy, and high-resolution structural analysis will be necessary to determine how spatially heterogeneous lacunar remodeling during reproduction contributes to the material behavior of maternal cortical bone.

Second, osteocyte relative age was inferred from fluorochrome mineral labeling and spatial location within the cortex, which provides indirect and relative information rather than precise temporal identification. Although this approach enabled the distinction between newly formed and pre-existing osteocytes, it did not allow the exact determination of osteocyte formation time. Genetic lineage-tracing models with an inducible fluorescent reporter may provide improved temporal resolution and enable more direct assessment of osteocyte age-related responses during reproduction. Third, the current analysis of lacunar remodeling was limited to the anterior and posterior cortices of the femoral midshaft. Although these midshaft cortical regions provide a well-defined model for studying spatial variation in lacunar remodeling, it remains unclear whether similar spatial patterns occur in other cortical regions, such as the medial and lateral cortices, or in skeletal sites with different loading environments and structural organization.

Extending this analysis to additional skeletal compartments, including the tibia and distal femur, will be important for determining the generalizability of these findings. Finally, the investigation of osteocyte cellular activity remains technically challenging due to their location within mineralized bone matrix. Conventional approaches based on cell isolation or *in vitro* culture do not preserve the native spatial and structural context of osteocytes, limiting the potential to interpret osteocyte activity in relation to their local microenvironment and spatial location within the cortex. Combining spatially resolved transcriptomic approaches with advanced imaging techniques in frozen bone sections may provide a more effective strategy for characterizing osteocyte signaling while preserving *in situ* morphology and spatial organization.

In summary, our study demonstrates that lacunar remodeling in maternal cortical bone during and after lactation is spatially heterogeneous. This spatial remodeling pattern is regulated by osteocyte PLR activity and varies with both osteocyte location and relative age. Newly formed osteocytes located near actively remodeling cortical surfaces were more responsive to the metabolic demands of reproduction, whereas pre-existing osteocytes embedded within cortical regions remained comparatively stable. In addition, osteocyte-level lacunar remodeling does not follow the same recovery pattern observed at the whole bone level after weaning. At the anterior endocortical region, lacunar enlargement during lactation was transient and largely resolved after weaning, whereas lacunar alterations at the posterior periosteal region persist beyond lactation.

Together, these findings demonstrate that skeletal recovery following lactation is not fully consistent across different levels of bone organization and identify osteocyte lacunar remodeling as a spatially organized and potentially persistent component of reproductive skeletal adaptation that may influence long-term maternal bone quality and skeletal health.

## 5. Acknowledge

Research reported in this publication was supported by the Penn Center for Musculoskeletal Diseases (PCMD) and NIH/NIAMS (P30-AR069619, R01-AR071718, and R56- AR071718).

## Supplementary Material

**Fig. S1.**
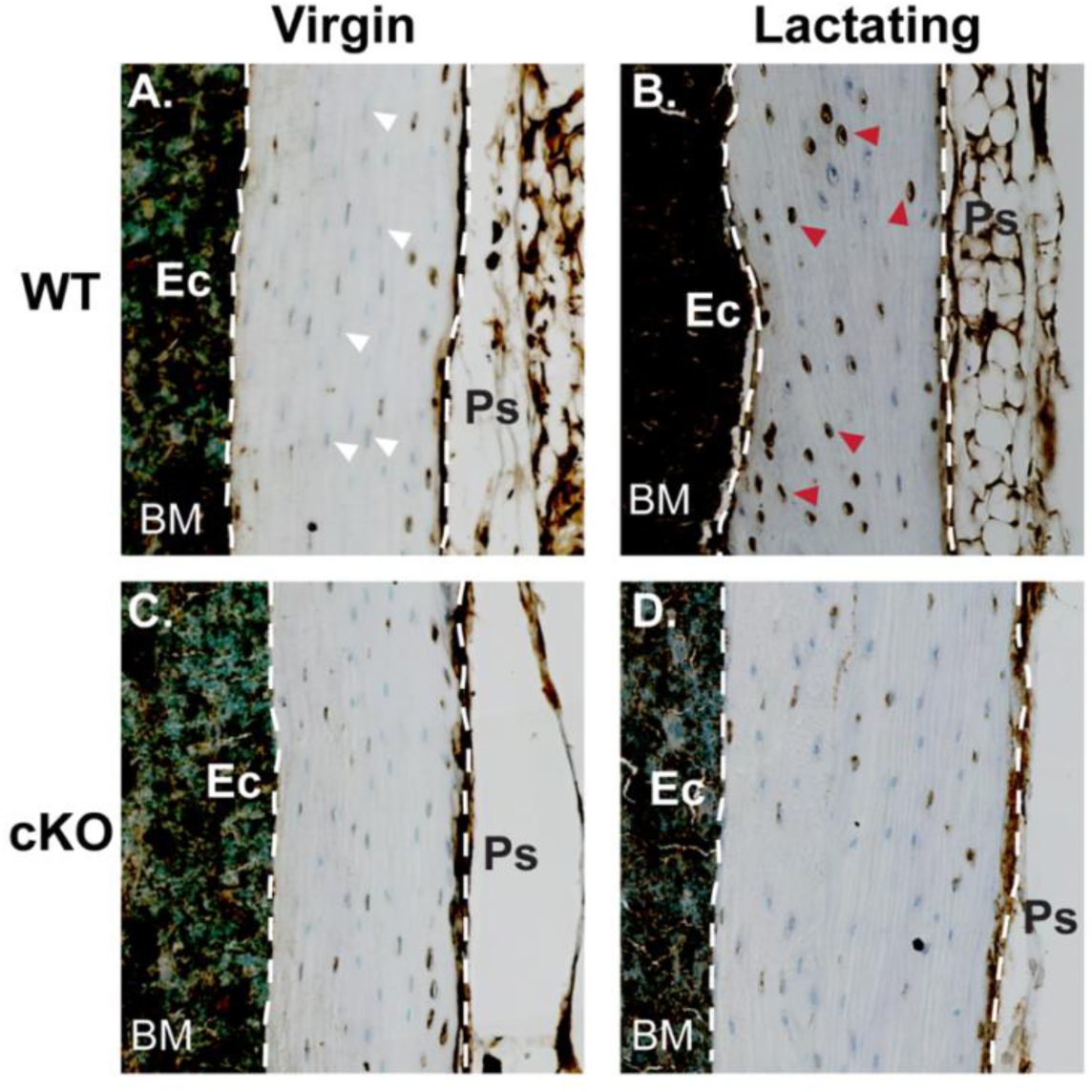
Representative immunohistochemistry (IHC) images of MMP13-stained osteocytes at the tibial midshaft in Virgin and Lactating WT (A, B) and cKO (C, D) mice. MMP13-positive osteocytes (brown) are indicated by red arrows (B), whereas MMP13-negative osteocytes with nuclear staining only (blue) are indicated by white arrows (A). Cortical boundaries are outlined by white dashed lines. Ps, periosteal surface; Ec, endosteal surface; BM, bone marrow.

**Fig. S2.**
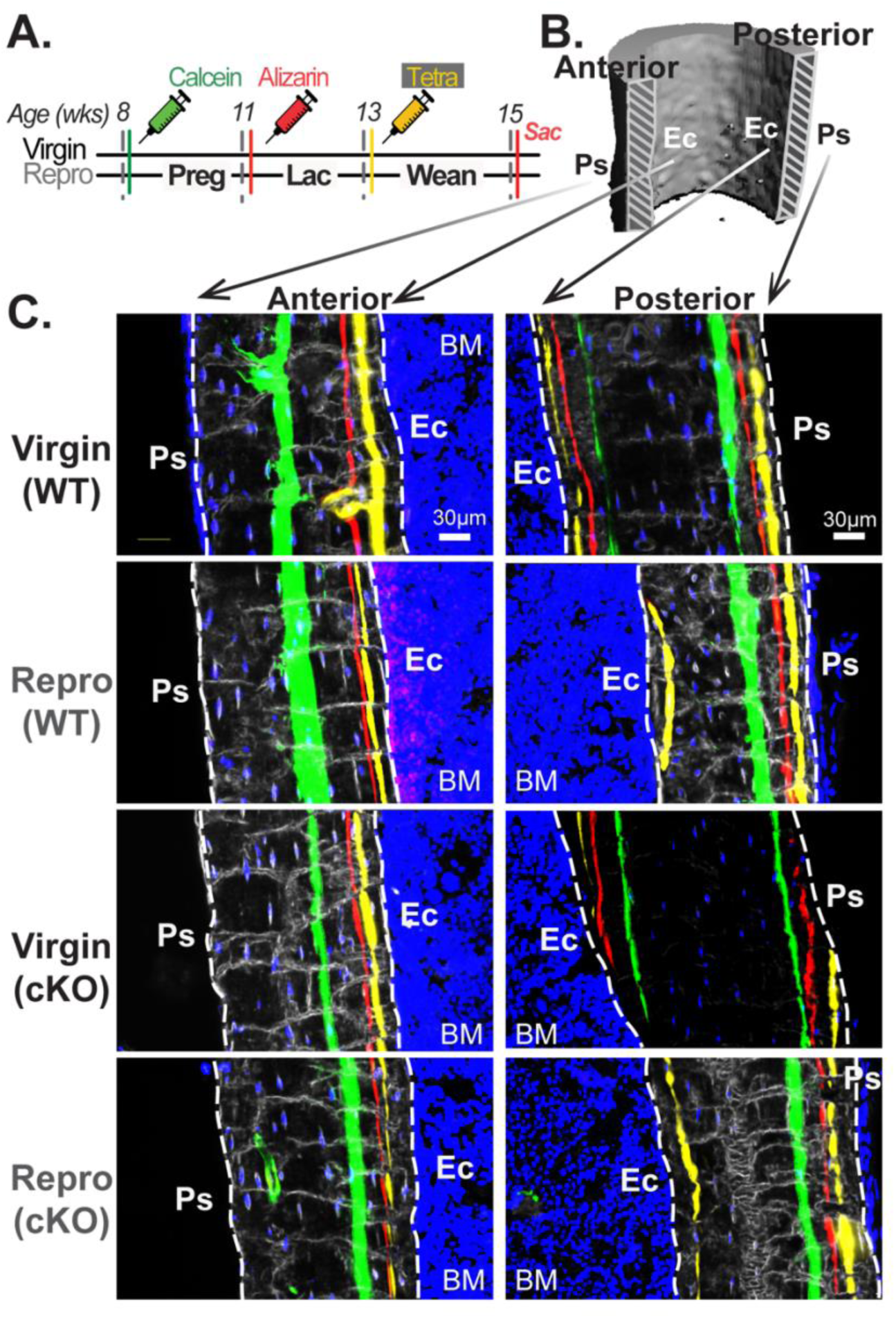
Fluorochrome labeling of dynamic mineral remodeling at the anterior and posterior cortices of the femoral midshaft in reproductive (Repro) and virgin WT and cKO mice. (A) Experimental timeline and fluorochrome labeling strategy. (B) Representative 3D reconstruction showing full cortical thickness from periosteal (Ps) to endosteal (Ec) surfaces at the femoral midshaft; grey-shaded regions indicate the anterior and posterior cortical areas analyzed. (C) Fluorochrome labeling at anterior (left) and posterior (right) cortices in Virgin and Repro WT and cKO mice. White dashed lines indicate cortical boundaries. Fluorescence labels: calcein (green), alizarin (red), tetracycline (Tetra; yellow), and To-PRO-3 (blue; nuclei). BM, bone marrow cavity. Scale bar: 30 μm.

**Fig. S3.**
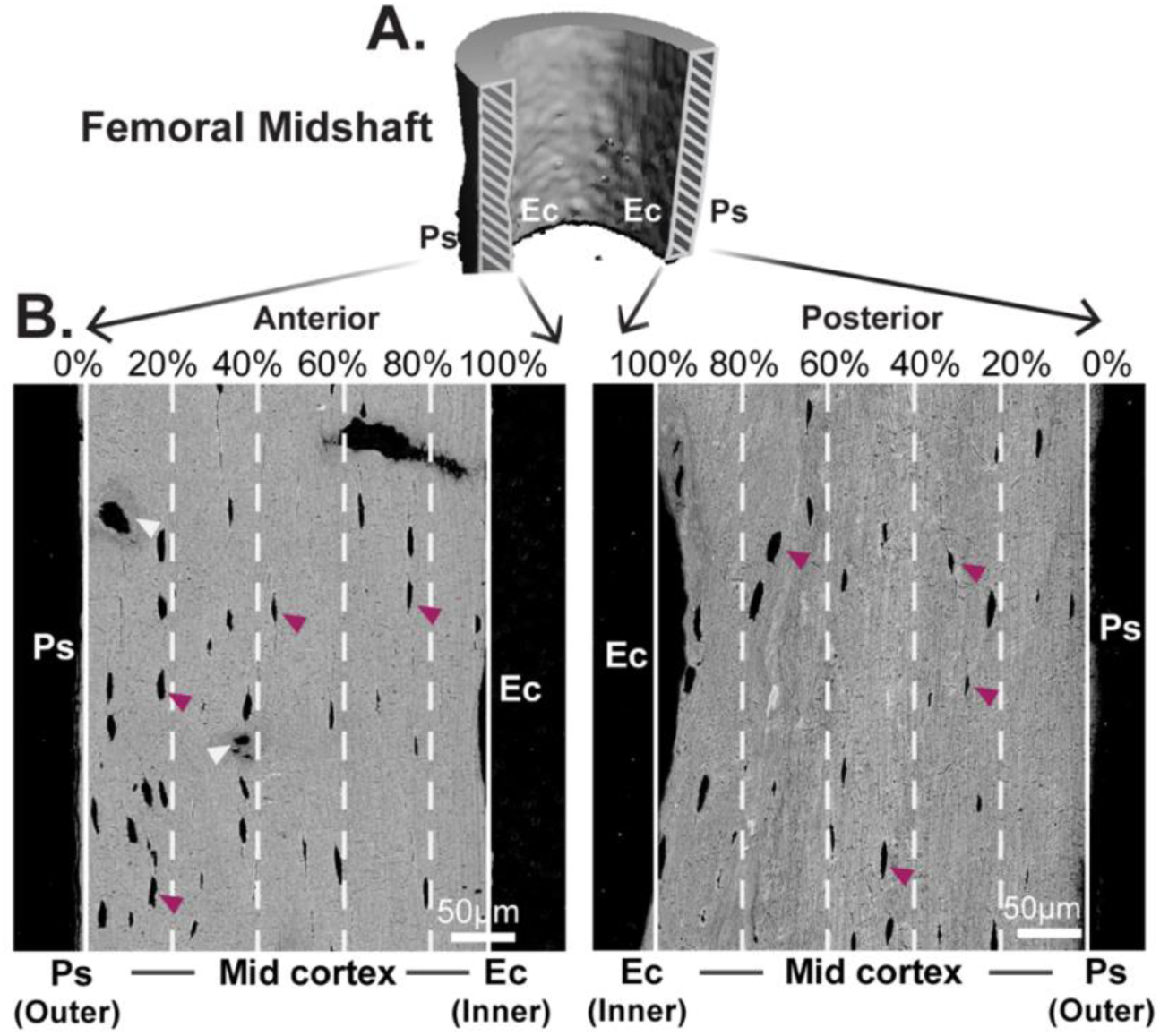
Representative bSEM images at the femoral midshaft for lacunar morphology analysis. (A) Representative 3D reconstruction of the longitudinal region of interest (ROI) at the anterior and posterior cortices, spanning the full cortical thickness from periosteal (Ps) to endosteal (Ec) surfaces. (B) Representative bSEM images of the anterior (left) and posterior (right) cortices in post-weaning WT mice, showing five equal sub-regions across the cortical thickness (0-100% from Ps to Ec; white dashed lines). Lacunar spaces are indicated by purple arrows, whereas larger voids marked by white arrows represent vascular spaces and were excluded from analysis. Scale bar: 50 μm.

**Table S1.**
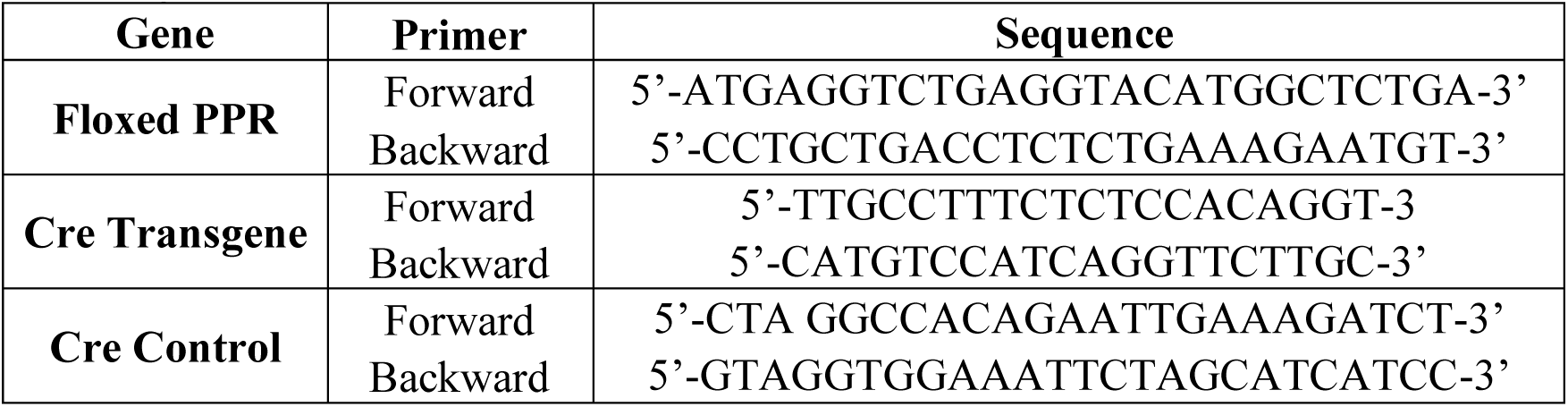
Primer information for genotyping by PCR.

**Table S2.** Animal information. (A) Body weight of Virgin and reproductive (Repro) mice with osteocyte-specific PPR deletion (cKO) and WT controls at each time point. (B) Litter size at parturition and average pup body weight at weaning in Repro cKO and WT mice. Differences in body weight relative to the pre-pregnancy baseline (8 weeks) were assessed within each group using paired Student’s t-tests (*p < 0.05).

| A.<br>Age\Group | Body Weight (g) |  |  |  |
| --- | --- | --- | --- | --- |
|  | Virgin |  | Reproduction |  |
|  | WT | cKO | WT | cKO |
| 8 wk (Pre-preg) | 15.4±0.3 | 16.6±1.3 | 17.7±0.7 | 14.9±1.0 |
| 11 wk (Preg) | 17.5±0.9* | 18.6±1.1 | 23.0±0.8* | 18.8±1.1* |
| 13 wk (Lac) | 17.2±0.7* | 19.1±1.0 | 25.4±2.8* | 21.4±2.5* |
| 15 wk (Wean) | 17.9±0.9* | 19.5±0.5 | 22.7±1.4* | 18.7±1.1* |

**Table S2. Animal information.** (A) Body weight of Virgin and reproductive (Repro) mice with osteocyte-specific PPR deletion (cKO) and WT controls at each time point.
| B.<br>Pups Information | Pups Information |  |
| --- | --- | --- |
|  | Reproductive group (Repro) |  |
|  | WT | cKO |
| Pups Num at Parturition | 5~7 | 6~7 |
| Pups Weight by Weaning (g) | 5.6±0.5 | 4.9±0.3 |

**Table S3.** Trabecular parameters measured by μCT in Virgin and reproductive (Repro) mice with osteocyte-specific PPR deletion (cKO) and WT controls at each time point. (A) Trabecular number (Tb.N), (B) trabecular separation (Tb.Sp), (C) connectivity density (Conn.D), and (D) structure model index (SMI). Differences relative to the pre-pregnancy baseline (8 weeks) were assessed within each group using paired Student’s t-tests (*p < 0.05). Differences relative to Virgin-WT at each time point were assessed using one-way ANOVA (+p < 0.05).

| A.<br>Age\Group | Tb.N |  |  |  |
| --- | --- | --- | --- | --- |
|  | Virgin |  | Repro |  |
|  | WT | cKO | WT | cKO |
| 8 wk (Pre-preg) | 3.00±0.18 | 3.50±0.40 | 3.98±0.65 | 3.81±0.49 |
| 11 wk (Preg) | 2.50±0.30* | 2.73±0.22* | 4.16±0.94+ | 3.73±0.33 |
| 13 wk (Lac) | 2.45±0.02* | 2.69±0.18 | 3.04±0.55 | 2.53±0.13* |
| 15 wk (Wean) | 2.26±0.30 | 2.27±0.23* | 2.44±0.24* | 2.49±0.22* |
| B.<br>Age\Group | Tb.Sp (mm) |  |  |  |
|  | Virgin |  | Repro |  |
|  | WT | cKO | WT | cKO |
| 8 wk (Pre-preg) | 0.34±0.02 | 0.30±0.04 | 0.26±0.05 | 0.27±0.03 |
| 11 wk (Preg) | 0.42±0.05* | 0.40±0.05 | 0.25±0.06+ | 0.27±0.03 |
| 13 wk (Lac) | 0.42±0.01* | 0.40±0.02 | 0.35±0.06+ | 0.40±0.02*+ |
| 15 wk (Wean) | 0.46±0.07 | 0.46±0.05* | 0.42±0.05* | 0.43±0.08* |
| C.<br>Age\Group | Conn.D (1/mm<sup>3</sup>) |  |  |  |
|  | Virgin |  | Repro |  |
|  | WT | cKO | WT | cKO |
| 8 wk (Pre-preg) | 21.80±10.25 | 23.52±11.80 | 54.56±17.10+ | 41.08±18.67 |
| 11 wk (Preg) | 19.47±2.31 | 12.30±4.61 | 80.70±49.96 | 49.28±12.68 |
| 13 wk (Lac) | 11.78±2.38 | 10.81±5.01 | 14.42±11.02* | 6.474±4.83* |
| 15 wk (Wean) | 10.29±1.88 | 7.14±5.23* | 15.43±4.40* | 12.13±2.63 |
| D.<br>Age\Group | SMI |  |  |  |
|  | Virgin |  | Repro |  |
|  | WT | cKO | WT | cKO |
| 8 wk (Pre-preg) | 3.08±0.19 | 3.17±0.17 | 2.82±0.18 | 3.04±0.26 |
| 11 wk (Preg) | 2.87±0.12 | 3.03±0.03 | 2.47±0.31* | 2.91±0.31 |
| 13 wk (Lac) | 3.069±0.10 | 3.07±0.15 | 3.44±0.26* | 3.70±0.28*+ |
| 15 wk (Wean) | 3.12±0.10 | 3.02±0.12 | 2.98±0.14 | 3.18±0.27 |

**Table S4.** Cortical parameters measured by μCT in Virgin and reproductive (Repro) mice with osteocyte-specific PPR deletion (cKO) and WT controls at each time point. (A) Cortical area (Ct.Area) and (B) polar Moment of Inertia (pMOI). Differences relative to the pre-pregnancy baseline (8 weeks) were assessed within each group using paired Student’s t-tests (*p < 0.05). Differences relative to Virgin-WT at each time point were assessed using one-way ANOVA (+p < 0.05).

| A.<br>Age\Group | Ct.Area (mm<sup>2</sup>) |  |  |  |
| --- | --- | --- | --- | --- |
|  | Virgin |  | Repro |  |
|  | WT | cKO | WT | cKO |
| 8 wk (Pre-preg) | 0.05±0.02 | 0.52±0.04 | 0.58±0.04 | 0.61±0.06+ |
| 11 wk (Preg) | 0.61±0.01* | 0.65±0.00* | 0.70±0.05*+ | 0.69±0.01+ |
| 13 wk (Lac) | 0.63±0.03* | 0.67±0.01* | 0.59±0.04* | 0.62±0.01 |
| 15 wk (Wean) | 0.65±0.03* | 0.70±0.02* | 0.67±0.06* | 0.67±0.03 |
| B.<br>Age\Group | pMOI (mm<sup>4</sup>) |  |  |  |
|  | Virgin |  | Repro |  |
|  | WT | cKO | WT | cKO |
| 8 wk (Pre-preg) | 0.18±0.01 | 0.20±0.02 | 0.23±0.02+ | 0.26±0.04+ |
| 11 wk (Preg) | 0.22±0.01* | 0.26±0.01* | 0.28±0.03*+ | 0.29±0.01+ |
| 13 wk (Lac) | 0.22±0.02* | 0.26±0.01* | 0.26±0.03* | 0.28±0.01+ |
| 15 wk (Wean) | 0.23±0.02* | 0.28±0.01* | 0.28±0.04* | 0.30±0.02+ |

